# A “torn bag mechanism” of small extracellular vesicle release via limiting membrane rupture of *en bloc* released amphisomes (amphiectosomes)

**DOI:** 10.1101/2024.01.30.578035

**Authors:** Tamás Visnovitz, Dorina Lenzinger, Anna Koncz, Péter M Vizi, Tünde Bárkai, Krisztina V Vukman, Alicia Galinsoga, Krisztina Németh, Kelsey Fletcher, Zsolt I Komlósi, Csaba Cserép, Ádám Dénes, Péter Lőrincz, Gábor Valcz, Edit I Buzás

**Affiliations:** Semmelweis University, Department of Genetics, Cell- and Immunobiology, Nagyvárad tér 4. 1089 Budapest, Hungary; ELTE Eötvös Loránd University, Department of Plant Physiology and Molecular Plant Biology, Pázmány Péter sétány 1/c, 1117 Budapest, Hungary; HUN-REN-SU Translational Extracellular Vesicle Research Group, Nagyvárad tér 4. 1089 Budapest, Hungary; Laboratory of Neuroimmunology, HUN-REN Institute of Experimental Medicine, Szigony u 43. 1083 Budapest, Hungary; ELTE Eötvös Loránd University, Department of Anatomy, Cell and Developmental Biology, Pázmány Péter sétány 1/c, 1117 Budapest, Hungary; Department of Image Analysis, 3DHISTECH Ltd, Budapest, Hungary; HCEMM-SU Extracellular Vesicle Research Group, Hungary, Nagyvárad tér 4. 1089 Budapest, Hungary

**Keywords:** exosomes, amphisome, extracellular vesicles, autophagy, secretion, biogenesis, migrasome

## Abstract

Recent studies showed an unexpected complexity of extracellular vesicle (EV) biogenesis pathways. We previously found evidence that human colorectal cancer cells *in vivo* release large multivesicular body-like structures *en bloc*. Here, we tested whether this large extracellular vesicle type is unique to colorectal cancer cells. We found that all cell types we studied (including different cell lines and cells in their original tissue environment) released multivesicular large EVs (MV-lEVs). We also demonstrated that upon spontaneous rupture of the limiting membrane of the MV-lEVs, their intraluminal vesicles (ILVs) escaped to the extracellular environment by a ”torn bag mechanism”. We proved that the MV-lEVs were released by ectocytosis of amphisomes (hence, we termed them amphiectosomes). Both ILVs of amphiectosomes and small EVs separated from conditioned media were either exclusively CD63 or LC3B positive. According to our model, upon fusion of multivesicular bodies with autophagosomes, fragments of the autophagosomal inner membrane curl up to form LC3B positive ILVs of amphisomes, while CD63 positive small EVs are of multivesicular body origin. Our data suggest a novel common release mechanism for small EVs, distinct from the exocytosis of multivesicular bodies or amphisomes, as well as the small ectosome release pathway.

## Introduction

Extracellular vesicles (EVs) are phospholipid bilayer enclosed structures^1–4^, which have important roles in cellular homeostasis and intercellular communication. Exosomes have been defined as small (~ 50-200 nm) EVs (sEVs) of endosomal origin^1,3,4^. Although autophagy is a major cellular homeostatic mechanism, and is implicated in a broad spectrum of human diseases, the intersection of autophagy and exosome secretion remains poorly understood. Recently, regulatory interactions have been shown between autophagy-related molecules and EV biogenesis^5,6^. Furthermore, the LC3-conjugation machinery was demonstrated to specify the cargo packaged into EVs^7^. Importantly, both others and we reported the secretion of LC3-carrying exosomes^7,8^. Particularly relevant to the findings presented here, is the implication of amphisomes (hybrid organelles formed by the fusion of late endosomes/multivesicular bodies (MVBs) with autophagosomes^9,10^) in EV biogenesis. It was suggested that fusion of the limiting membrane of amphisomes with the plasma membrane of cells results in a subsequent release of exosomes by exocytosis^1,3,11^. The current study was prompted by our recent data showing the *in vivo en bloc* release of large, MVB-like sEV clusters by human colorectal cancer cells^12^. Here we investigated if this was a colorectal cancer cell-specific phenomenon. Unexpectedly, we found that it was a general mechanism of sEV release that we designated as “torn bag mechanism”.

## Results and Discussion

In this study, we analyzed *in situ* fixed, cultured cells with the released EVs preserved in their original microenvironment on a surface coated by gelatin and fibronectin. We detected large multivesicular EVs (MV-lEVs) in sections of different immersion fixed organs. We tested tumorous HT29, HepG2, and non-tumorous HEK293, HEK293T-PalmGFP, HL1 cell lines, as well as primary suspension type bone marrow derived mast cells (BMMCs). In addition, we studied ultrathin sections of mouse kidney and liver.

By the analysis of transmission electron micrographs of all tested cell types, we identified budding (Fig.1A-G) and secretion (Fig.1H-N) of MV-lEVs carrying intraluminal vesicles (ILVs). Importantly, in all cases we found evidence for the extracellular rupture of the limiting membrane of MV-lEVs and the release of ILVs (Fig.1O-U). For this novel type of sEV release, we suggest the designation “torn bag mechanism”, which is distinct from the exocytosis of MVBs and amphisomes^1,3,4,11^ and from the release of plasma membrane derived small EVs (sEVs) by ectocytosis^13^.

**Figure 1.**
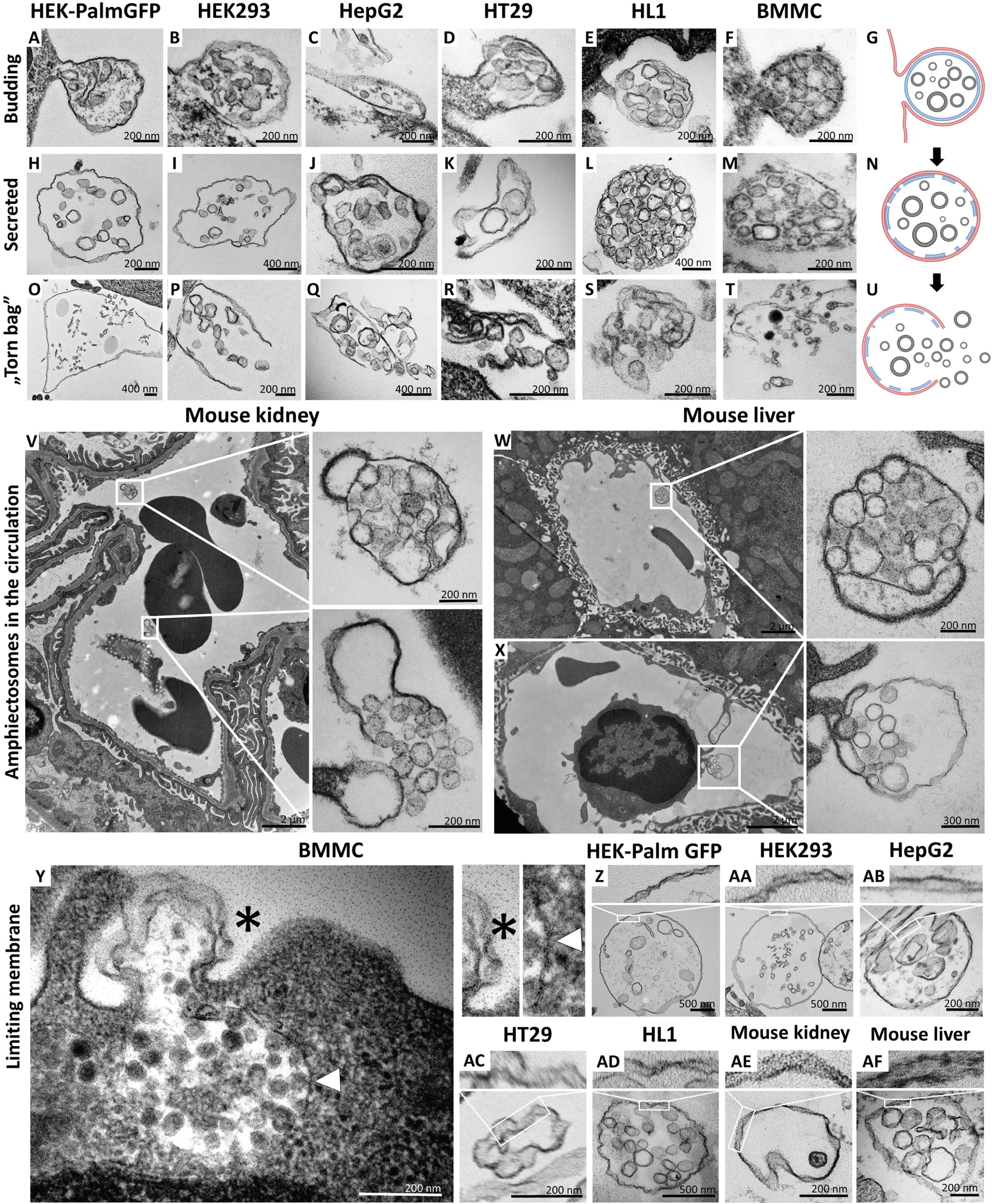
Transmission electron microscopic detection of the release and extracellular fate of large, multivesicular EVs secreted by different cell lines and cells in mouse organs. Major steps of the release of large, multivesicular EVs (MV-lEVs) were detected in the case of all tested cell lines including the immortal, non-tumorous HEK293T-PalmGFP (A,H,O), HEK293 (B,I,P), the tumorous cell lines HepG2 (C,J,Q) and HT29 (D,K,R), the beating cardiomyocyte cell line HL1 (E,L,S) and the primary suspension of bone marrow derived mast cells (BMMCs) (F,M,T). The different phases of EV secretion were also captured in the circulation of mouse kidney (V) and liver (W,X). According to the electron micrographs, we found evidence for the budding (A-G,X) and secretion (H-N,V,W) of the MV-lEVs. We also detected the extracellular rupture of the limiting membrane of the released MV-lEVs with the escape of the intraluminal vesicles (ILVs) by a “torn bag mechanism” (O-U,V). Although it is not always clear whether the secreted MV-lEVs have a single or double limiting membrane, several micrographs suggest the presence of the double membrane (Y-AF) in the secreted MV-lEVs. In the case of BMMCs (Y), the release phase of a multivesicular structure is captured. The bottom portion of this structure embedded in the cytoplasm is surrounded by a single membrane (white arrow head) while the upper (budding) portion is covered by double membrane (asterisk). In the shematic figures (G, N, U) the limiting membrane of MV-lEV presumably with plasma-membrane origin was indicated by red, the original limiting membrane of intracellular amphisomes, which may fragmented during the release process was indicated by blue while the ILVs of the MV-lEV were shown by gray color.

Most relevant to the *in vivo* conditions, we also observed the same phenomenon within the ultrathin sections of both murine kidney (Fig.1V) and liver (Fig.1W,X). In these cases, both the intact MV-lEVs (Fig.1V-X) and the “torn bag release” of sEVs (Fig.1V) were detected. Fig.1X shows that a circulating leukocyte releases MV-lEVs by ectocytosis. In Fig.1_S1, MV-lEVs, lEVs and sEVs were captured simultaneously both in kidney (Fig.1_S1C,D) and liver (Fig.1_S1F,H,I). In the mouse liver section (Fig.1_S1G), MV-lEV secretion both by endothelial and subendothelial cells can be detected.

Based on the transmission electron microscopy (TEM) analysis of ultrathin sections, it was not always obvious whether the secreted MV-lEVs had a single or double membrane. However, several micrographs suggested an at least partially intact double membrane (Fig.1Y-AF) of MV-lEVs. In the case of BMMCs (Fig.1Y), the release phase of a multivesicular structure is captured. The bottom portion of this structure, embedded in the cytoplasm, is surrounded by a single membrane while the upper (budding) portion is covered by double membrane. We hypothesize that disruption of the original amphisome membrane mainly occurs after separation of the MV-lEV from the cell to avoid the release of ILVs inside the cell.

Next, we decided to further investigate the subcellular origin of the ILVs within the secreted MV-lEVs. First, we analyzed the microenvironment of *in situ* fixed HEK293T-PalmGFP cells by confocal microscopy. The PalmGFP signal of HEK293T-PalmGFP cells principally associates with the plasma membrane^14^ (Fig.2_S1A-C), therefore the green fluorescence helped us to identify the plasma membrane-derived limiting membrane of MV-lEVs. In agreement with our previous findings on HT29 colorectal cancer cells, within the MV-lEVs, we found CD63/ALIX (Fig.2A,G), CD81/ALIX (Fig.2B,H), CD63/TSG101 (Fig.2.C,I) and CD81/TSG101 (Fig.2D,J) double positive ILVs or ILV clusters.

**Figure 2.**
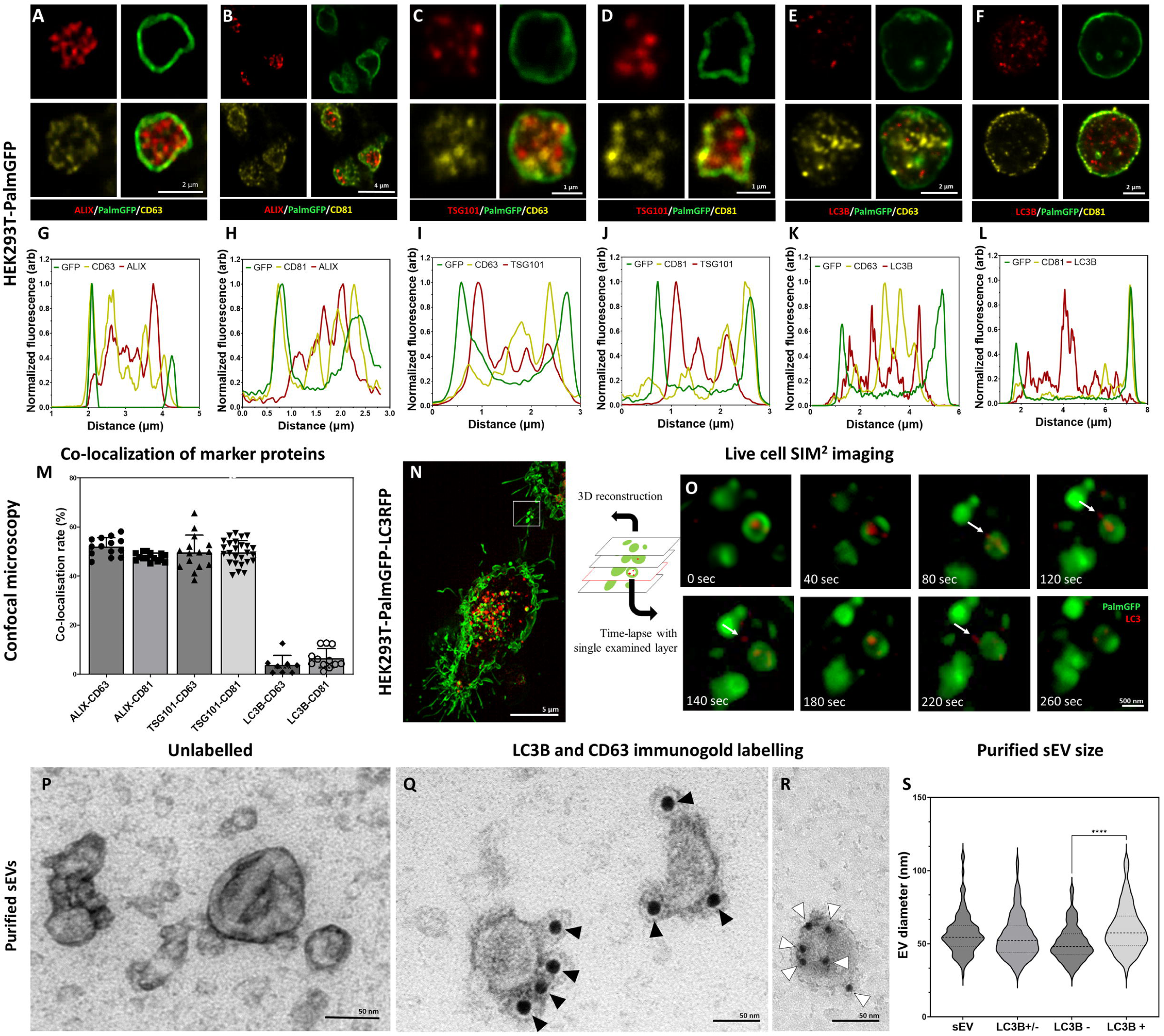
Detection of conventional sEV markers and the LC3 protein in HEK293T-PalmGFP cell-derived EVs. Widely used sEV markers (CD63, CD81, ALIX and TSG101) and LC3B were tested in MV-lEVs found in the microenvironment of the releasing cells by confocal microscopy after *in situ* fixation (A-F). Normalized fluorescence intensities were calculated in order to determine the relative localization of the limiting membrane (PalmGFP), the conventional sEV markers and the LC3B signal (G-L). Fluorescence intensity peaks of sEV markers were largely overlapping with each other, while the LC3B signal and the sEV markers showed separation. Co-localization rates were also calculated (M). The sEV markers co-localized with one another as no significant difference was found among them. In contrast, low co-localization rates were detected between the “classical” sEV markers and LC3B (one-way ANOVA, p<0.0001, n=8-26 confocal images). Real time release of LC3 positive sEVs by the “torn bag mechanism” was studied in the case of HEK293T-PalmGFP-LC3RFP cells by Elyra7 SIM^2^ super-resolution live cell imaging (N,O). Images were recorded continuously, and selected serial time points are shown. LC3 positive, red fluorescent small particles were released within a 5 min timeframe (O) and are indicated by white arrows. Presence of CD63 and LC3B were detected in the case of a sEV fraction separated from serum-free condition medium using immunogold transmission electronmicroscopy (TEM). HEK293T-PalmGFP derived sEV fraction is shown by negative-positive contrast without immune labelling (P). In double-labelled immunogold TEM images (Q,R), distinct LC3B positive (Q) and CD63 positive (R) sEVs were found. However, CD63-LC3B double positive EVs were not detected. Black arrowheads indicate 10 nm gold particles identifying LC3B, while white arrowheads show 5 nm gold particles corresponding to the presence of CD63. Quantitative analysis of TEM images was performed (S), and the diameters of different EV populations were determined. The LC3B negative population was significantly smaller than the LC3B positive one (p<0.0001, t-test; n=79-100). No difference was detected when the immunogold labelled sEV fraction (either LC3B positive or negative, LC3B+/-) and the unlabeled sEV fraction (sEV) were compared (p<0.05, t-test, n=112-179).

We also studied the possible autophagy related aspects of the secreted MV-lEVs. ILVs were tested for the autophagy marker LC3B in parallel with CD63 and CD81. Although LC3B, CD63 and CD81 were all present in association with the ILVs (Fig.2E,F), the LC3B and CD63 (Fig.2K) and the LC3B and CD81 (Fig.2L) signals did not overlap. Fig.2M shows that while the known sEV markers (CD63, CD81, TSG101 and ALIX) strongly co-localized with each other, LC3B positivity hardly showed colocalization with CD63 or CD81. Immunocytochemistry analysis of HT29, HepG2 and the cardiomyoblast H9c2 cells further validated the findings obtained with the HEK293T-PalmGFP cells (Fig.2_S2 and Fig.2_S3A,B). The ILVs of HEK293T-PalmGFP and HepG2 cell lines were also Rab7 positive (Fig.2_S3C,D), suggesting a late endosomal origin. Western blotting of the applied antibodies is summarized in Fig.2_S4.

The sEV markers were also tested by TEM using negative-positive contrasting technique^15^ on sEVs separated form serum-free conditioned medium of HEK293T-PalmGFP cells. Fig.2P confirms the typical sEV morphology. With TEM double immunogold labelling, using anti-LC3B and anti-CD63 antibodies simultaneously, we found distinct LC3B positive (Fig.2Q, Fig.2_S6O) and CD63 positive (Fig.2R, Fig.2_S6O) sEVs. Based on the analysis of TEM images, the diameters of unlabeled, and LC3B positive and negative sEVs were determined (Fig.2S). The LC3B positive sEVs had a significantly larger diameter as compared to the LC3B negative ones.

To conclude our marker studies, we detected the presence of CD63, CD81, TSG101, ALIX positive, most probably MVB-derived ILVs. In addition, the autophagosome marker carrying, LC3B positive ILVs were also found within the same single, plasma membrane limited extracellular MV-lEV, which identified these MV-lEVs as *en bloc* released amphisomes^16^ that we refer to “amphiectosomes”.

The “torn bag mechanism” was also monitored by live cell SIM^2^ super-resolution microscopy analysis of HEK293T-PalmGFP-LC3RFP cells (Fig.2N,O). The release of the LC3 positive red fluorescent signal was detected within a relatively short period of time (the first LC3 positive ILVs left the amphiectosome within 40s, the whole “torn bag” sEV release process was completed within 260 sec) (Fig.2O). We could rule out the possibility that rupture of the limiting membrane detected by TEM (Fig.1O-T,V) was a fixation artefact by showing the spontaneous release of LC3 positive sEVs from amphiectosomes with live-cell imaging. Characterization of the in-house developed HEK293T-PalmGFP-LC3RFP cell line is shown in Fig.2_S5.

In the following step, we addressed the question whether LC3, associated with the ILVs of MV-lEVs, indeed reflected autophagy origin. We tested MVBs (Fig.2_S6A,F,K), autophagosomes (Fig.2_S6B,G,L), amphisomes (Fig.2_S6C,H,M), amphiectosomes (Fig.2_S6D,I,N) and isolated sEV fractions of the same cells (Fig.2_S6E,J,O). Using immune electron microscopy, as expected, we found CD63 single positivity in MVBs (Fig.2_S6K). In autophagosomes, the limiting phagophore membrane was LC3B positive, and CD63 positivity was also present (Fig.2_S6L). The limiting membrane of amphisomes was LC3B negative, and the internal membranous structures were either LC3B or CD63 positive (Fig.2_S6M). The same immunoreactivity was also observed in the ILVs of the released amphiectosomes (Fig.2_S6N). Importantly, sEVs separated form serum-free conditioned medium of HEK293T-PalmGFP cells were either LC3B or CD63 positive (Fig.2_S6O). Thus, we confirmed our confocal microscopy results at the ultrastructural level. Using immunogold TEM, we provided further evidence for the budding/ectocytosis mechanism of amphiectosome release (Fig.2_S6P). The diameters of ILVs within MVBs, amphisomes and amphiectosomes were compared (Fig.2_S6Q), and the differences were likely due to the different membrane composition, pH and osmotic conditions within these structures. In agreement with our observations with separated sEVs, LC3B positive ILVs had a significantly larger diameter than the LC3B negative ones (Fig.2_S6R) possibly indicating difference in membrane composition and their different intracellular origin. Based on all the above findings, we propose the following model (Fig.3A): autophagosomes and MVBs fuse to form amphisomes, and the inner, LC3 positive membrane of autophagosomes undergoes fragmentation^16^. Membrane fragments curl up and form LC3 positive ILVs. Therefore, amphisomes contain both MVB-derived CD63 positive/LC3 negative and autophagosome-derived, CD63 negative/LC3 positive ILVs. The amphisome is next released from the cell by ectocytosis. Finally, the plasma membrane-derived outer membrane ruptures enabling the ILVs escape to the extracellular space by a “torn bag mechanism”. By using stimulated emission depletion (STED) microscopy, we documented the intracellular phases of our proposed model: MVB (Fig3.B), autophagosome (Fig3.D), the fusion of MVB and autophagosome (Fig3.C), fragmentation of the LC3 positive membrane and ILV formation from the membrane fragments (Fig3.F) and mature amphisome (Fig.3E). The plasma membrane origin of the external membrane of amphiectosome was further supported by wheat germ agglutinin-based live cell labelling (Fig.3G).

**Figure 3.**
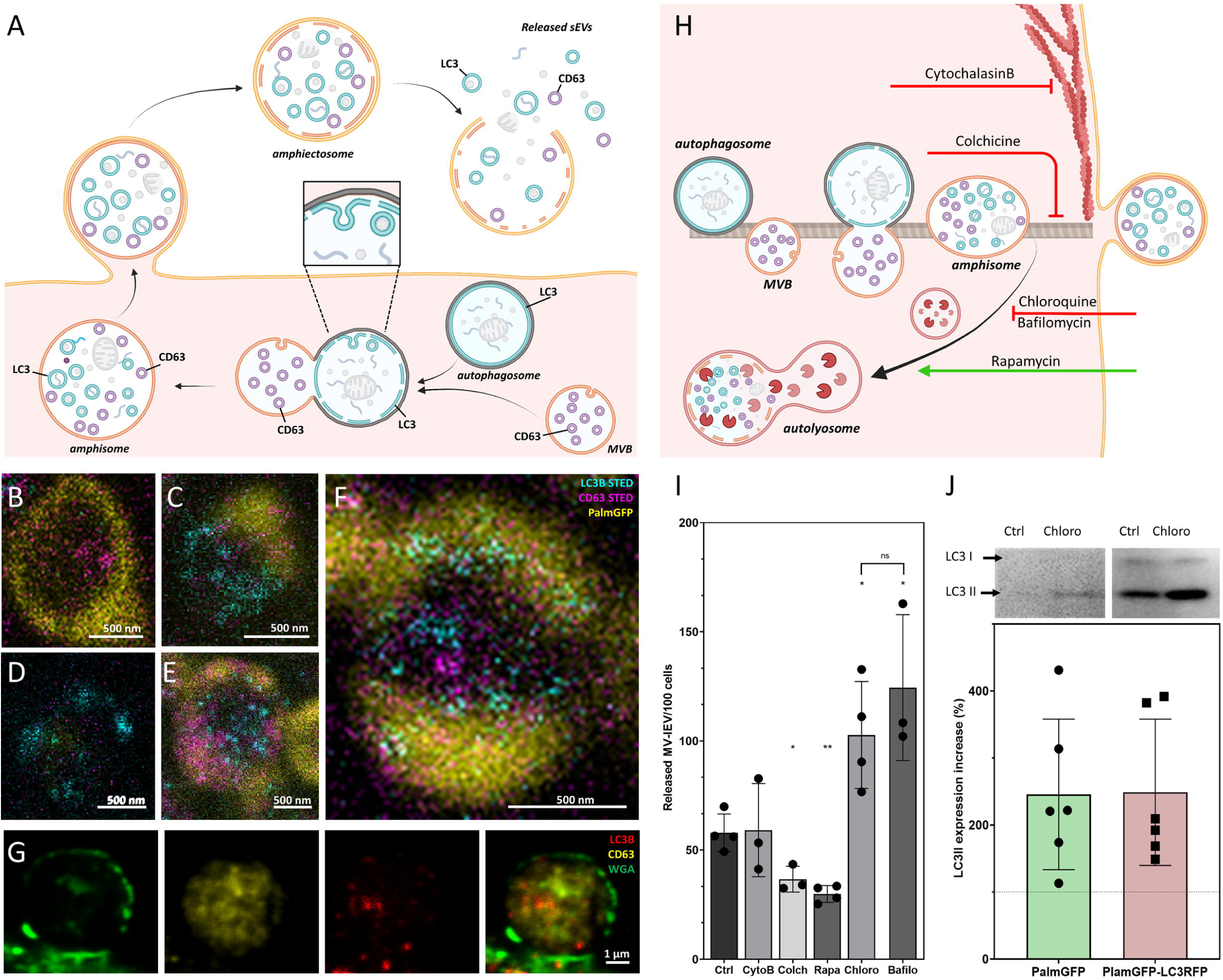
Amphiectosome release and its modulation. Based on our data, a model of amphiectosome release was generated (A). According to this model, the fusion of MVBs and autophagosomes forms amphisomes. The LC3B positive membrane layer (indicated in cyan) undergoes disintegration and forms LC3B positive ILVs inside the amphisome. Later, the amphisome is released into the extracellular space by ectocytosis and can be identified extracellularly as an amphiectosome. Finally, the limiting membrane(s) of the amphiectosome is ruptured and the ILVs are released as sEVs into the extracellular space by a “torn bag mechanism”. Steps of amphisome formation including LC3 positive ILV formation in 30 µM Chloroquine-treated HEK293T-PalmGFP cells was followed by super-resolution (STED) microscopy (B-F). The super-resolution STED channels were LC3B (cyan) and CD63 (magenta), while yellow indicates the confocal PalmGFP signal. Intracellular vesicular structures (such as endosomes, MVBs and amphisomes) may recieve Palm-GFP from the plasma membrane. An MVB (B), an autophagosome with Palm-GFP negative membrane (D), fusion of an autophagosome and an MVB (C), formation of LC3B positive ILVs in an amphisome (F) and a mature amphisome (E) were detected. To confirm the origin of the external membrane layer of amphiectosomes, fluorescently labelled WGA was applied. The plasma membrane of the living non-fluorescent HEK293 cells was labelled. As the external membrane of the budding amphiectosome was WGA positive, its plasma membrane origin is confirmed (G). In order to further support our model on amphiectosome release and “torn bag” EV secretion, different *in vitro* treatments were applied. Cytochalasin B, Colchicine, Chloroquine, Bafilomycin A1 and Rapamycin were used to modulate amphiectosome release. Targeted molecular processes are summarized (H). While Cytochalasin B inhibits actin-dependent membrane budding and cell migration, Colchicine blocks the microtubule-dependent intracellular trafficking. While Chloroquine and Bafilomycin have similar, Rapamycin has opposite effect on lysosome-autophagosome or lysosome-amphisome fusion. Chloroquine and Bafilomycin inhibit lysosomal degradation while Rapamycin accelerates it. Based on confocal microscopy, Cytochalasin B (CytoB) did not alter the dynamics of amphiectosome release (I). In contrast, both Colchicine (Colch) and Rapamycin (Rapa) significantly inhibited the release of amphiectosomes, while Chloroquine (Chloro) and Bafilomycin (Bafilo) increased the release frequency. There was no difference between the effect of Chloroquine and Bafilomycin (I). Results are shown as mean ± SD of 3-4 independent biological replicates, analyzed by one-way ANOVA and Student’s t test, *: p<0.05, **: p<0.01, ns: non-significant. Original LASX files, which served as a basis of our quantification, are publicly available (doi: 10.6019/S-BIAD1456). Example for the calculation is shown on Fig3_S1H. Presence of membrane-bound (lipidated) LC3II was tested by Western blotting. The total protein content of serum-, cell- and large EV-depleted conditioned medium of HEK293T-PalmGFP (PalmGFP) and HEK293T-PalmGFP-LC3RFP (PalmGFP-LC3RFP) cells was precipitated by TCA and 20 µg of the protein samples were loaded on the gel (J). The lipidated LC3II band was detected in all cases. Relative expression of control (Ctrl) and Chloroquine (Chloro)-treated samples were determined by densitometry. Chloroquine treatment increased the LC3II level by approximately two fold. Results are shown as mean ± SD of n=6 biological replicates.

To investigate the process of amphiectosome release, we exposed the MV-lEV releasing cells to different *in vitro* treatments (Fig.3H). The release of MV-lEVs was monitored by confocal microscopy of *in situ* fixed cell cultures. Optimal test conditions were determined (Fig.3_S1A-F) and the results are summarized in Fig.3I. Original LASX files which served as a basis of our quantification, are publicly available (doi: 10.6019/S-BIAD1456). An example for our approach of counting the MV-lEVs is shown on Fig3_S1H. Cytochalasin B did not have any effect on the discharge of MV-lEVs suggesting that the release did not involve a major actin-dependent mechanism. In contrast, there was a significant reduction of the MV-lEV secretion upon exposure of the cells to Colchicine indicating a role of microtubules in the release of the MV-lEVs. While Rapamycin significantly reduced the discharge of MV-lEVs, Chloroquine and Bafilomycin induced an enhanced MV-lEV secretion. Rapamycin activates autophagic degradation^17^, therefore, it induces a shift towards degradation as opposed to secretion. The lysosomotropic agents Chloroquine and Bafilomycin are known to interfere with the acidification of lysosomes^18,19^. By blocking the degradation pathway of MVBs/amphisomes (Fig.3H), an enhanced sEV secretion is observed. This effect is well known for exosome secretion from MVBs^20,21^. The diameters of the released MV-lEVs were determined based on confocal images (Fig.3_S1G). Metabolic activity of the cells was determined by a Resazurin assay, and a significant reduction was detected upon exposure of the cells to Rapamycin (Fig.3_S1D) in line with previously published data^22^. LC3II is the membrane-associated, lipidated autophagic form of LC3^23^ and it is the hallmark of autophagy related membranes^16^. Importantly, by Western blot, we not only showed the presence of the membrane-bound LC3II in serum-free, lEV-depleted (sEV containing) conditioned medium of both HEK293T-PalmGFP and HEK293T-PalmGFP-LC3RFP cells, but the amount of LC3II substantially increased upon Chloroquine treatment (Fig.3J). Raw data of Western blots are available in Fig.3_S2.

Recent advances in the EV field shed light on migrasomes, a special type of MV-lEVs^24,25^. With their pomegranate-like ultrastructure, migrasomes resemble amphiectosomes. Therefore, we tested the presence of TSPAN4, a migrasome limiting membrane marker^25^, in amphiectosomes. Fig.4A,B,G,H show that although TSPAN4 was present intraluminally in the HEK293T-PalmGFP-derived MV-lEVs, it was clearly absent from their external membrane. Surprisingly, we identified two different HT29 cell-derived MV-lEV populations: one in which TSPAN4 was only located intraluminally (Fig.4C,E,I,K), and another one with a TSPAN4 positive external membrane (Fig.4D,F,J,L). This raised the possibility that the latter population corresponded to migrasomes. Our co-localization analysis also confirmed the existence of two distinct MV-lEV populations (Fig.4M). Next, we carried out live-cell imaging on HEK293T-PalmGFP-LC3RFP cells. The released MV-lEVs were either LC3 positive or negative intraluminally (Fig.4N,O). Our TEM images confirmed that certain cell types can release both migrasome-like structures and amphiectosomes. MV-lEVs with typical migrasome-associated retraction fiber(s) were detected in the case of HL1 (Fig.4P), HEK293T-PalmGFP (Fig.4Q) and BMMC cells (Fig.4R). Of note, it cannot be excluded that the elongated structures observed in the above cases, may correspond to tunnelling nanotubes^26^. Importantly, the same cell lines also released amphiectosomes by budding from the cell surface (Fig.4S-U). Taken together, based on the absence of TSPAN4 in their external membrane, and their lack of association with retraction fibers, amphiectosomes appear to be distinct from migrasomes. Besides migrasomes, another MV-lEV type was described in the case of gastrointestinal tumors and low-grade glioblastoma cells referred to as spheresome^27,28^. However, there is no data on a relationship of spheresome release and autophagy. Recently, endothelial cell derived, multicompartmented microvesicles (MCMVs) were shown to protrude and pinch-off from the cell surface releasing ILVs by a mechanism similar to exocytosis^29^. The absence of protrusion clusters described for MCMV^29^ distinguishes amphiectosomes from MCMVs. In addition, the previously decribed so called “nodal vesicular parcels”^30^ might be special examples of amphiectosomes. Finally, in C. elegans, the release autophagy and stress related large EVs (exophers) has been documented^31,32,33^. They contain damaged organelles and do not have a MV-lEV like ultrastructure. In contrast, the amphiectosomes we described here, have multivesicular structure without recognizable damaged organelles.

**Figure 4.**
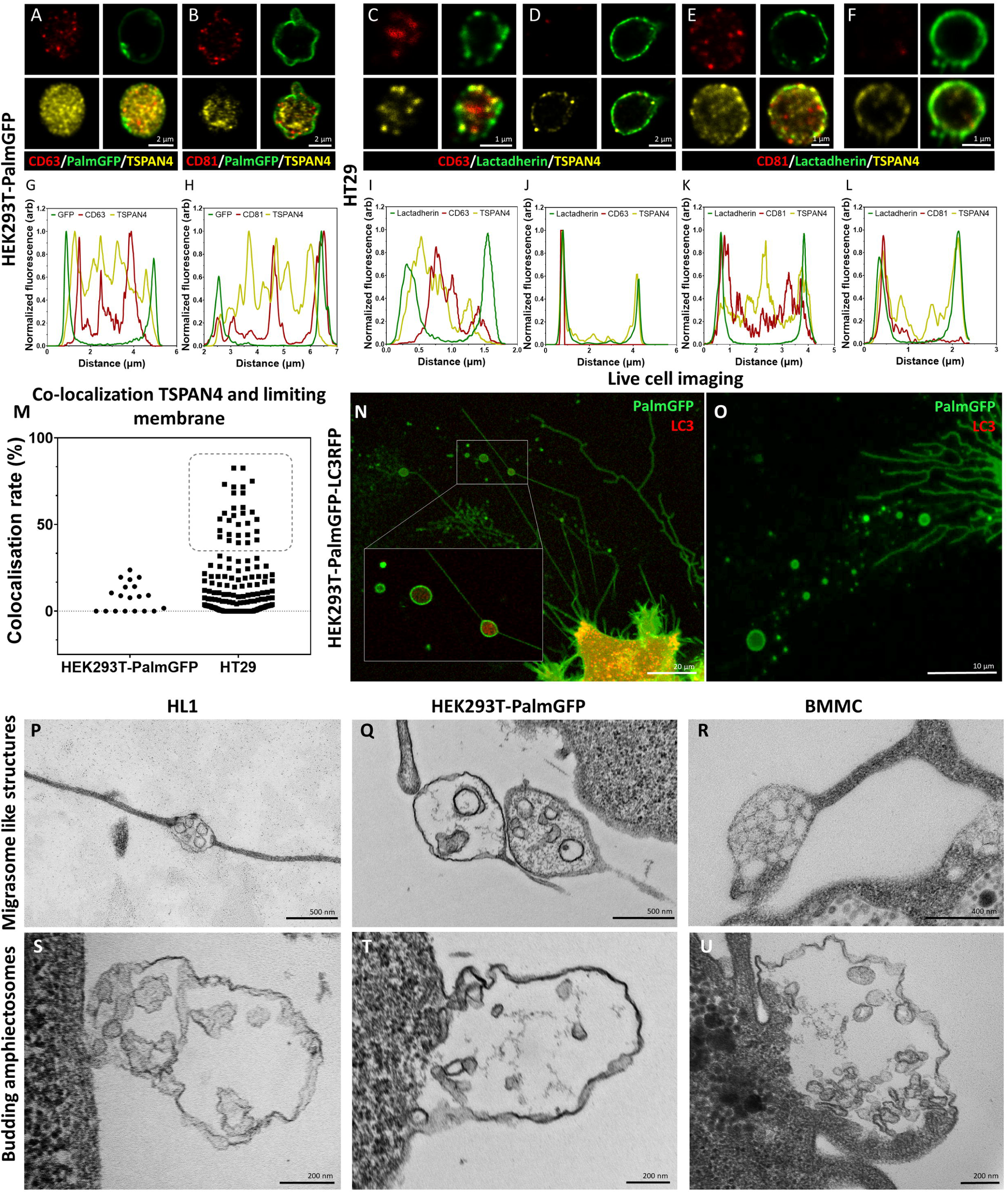
Comparison of amphiectosomes and migrasomes. Commonly used sEV markers (CD63, CD81) and TSPAN4, a suggested migrasome marker, were tested in *in situ* fixed intact MV-lEVs of HEK293T-PalmGFP (A,B) and HT29 (C-F) cells by confocal microscopy. Normalized fluorescence intensities were calculated in order to determine the relative localization of the limiting membrane (with PalmGFP or lactadherin labelling) and the CD63/TSPAN4 and CD81/TSPAN4 markers (G-L). In the case of HEK293T-PalmGFP-derived EVs, we did not find migrasomes with TSPAN4 in their limiting membrane. The TSPAN4 signal was only detected intraluminally in the MV-lEVs. The limiting membranes of HT29-derived MV-lEVs were either TSPAN4 positive and negative. The co-localization rate between the limiting membrane and TSPAN4 was low in case of HEK293T-PalmGFP-derived EVs. In the case of HT29 cells, two MV-lEV populations were identified: one with low and one with high co-localization rates (M). Live cell imaging of HEK293T-PalmGFP-LC3RFP cells showed retraction fiberassociated MV-lEVs with or without intraluminal LC3 positivity (N,O). Using TEM, we could identify structures with retraction fiber-associated morphology in the case of HL1 (P), HEK293T-PalmGFP (Q) and BMMC (R) cells. For comparison, budding of amphiectosomes of the same HL1 (S), HEK293TPalmGFP (T) and BMMC cells (U) are shown (without being associated with long retractions fibers).

Our approach, involving *in situ* fixation of cultures and tissues, made it possible to recognize sEV release from amphiectosomes by the “torn bag mechanism”. We propose that this mechanism could be easily missed earlier if conditioned medium was subjected to centrifugation, SEC purification or even to simple pipetting, which may rupture the limiting membrane of amphiectosomes. This aligns with our observation that the spontaneous escape of ILVs from untouched amphiectosomes can be completed as early as 5 minutes after amphiectosome release. Based on our data presented here, and considering that the exocytosis of MVBs/amphisomes under steady-state conditions is rarely documented in the scientific literature, we suggest that amphiectosome secretion and the “torn bag mechanism” may have a significant, yet previously unrecognized, role in sEV biogenesis.

## Material and methods

### Key Resources Table

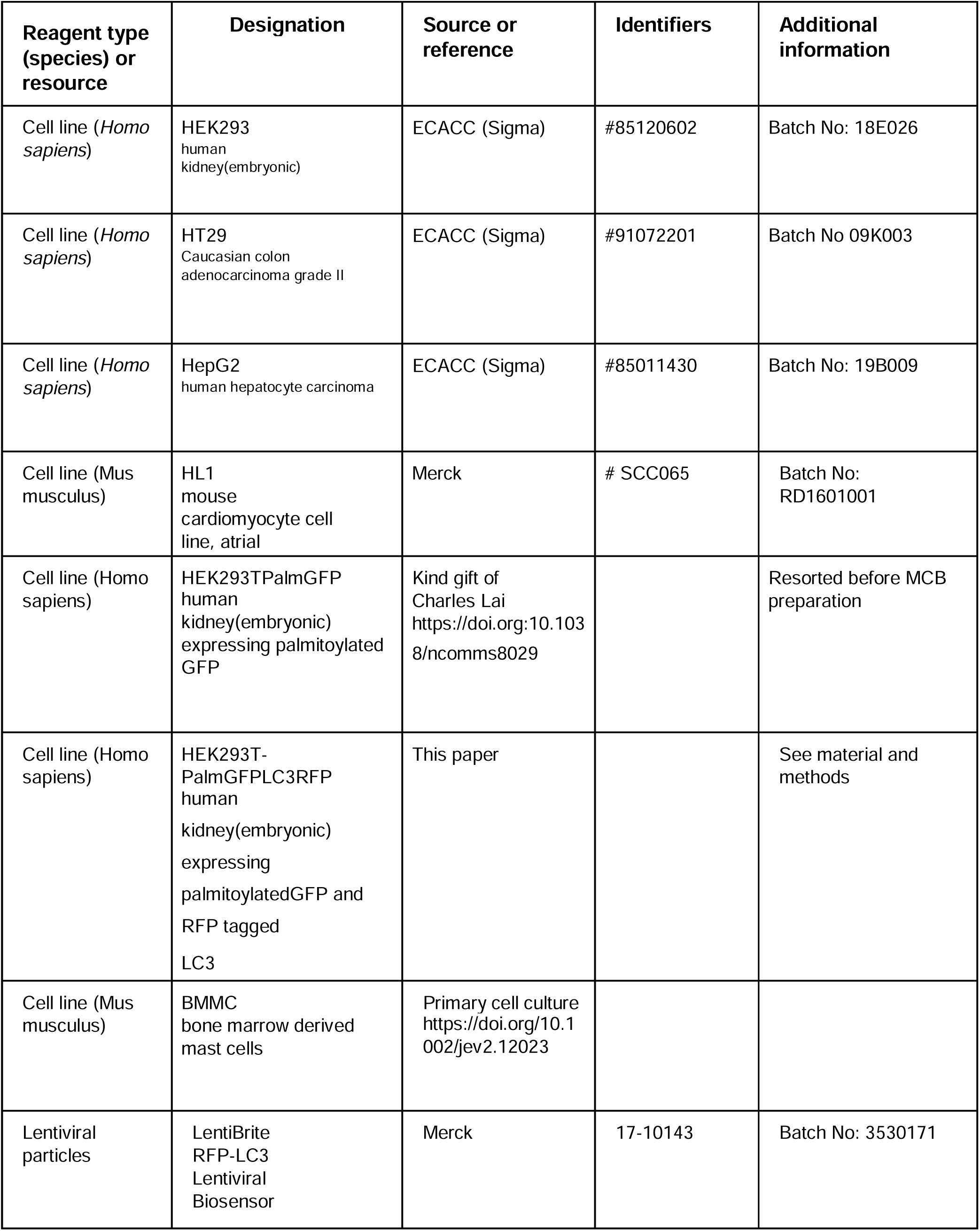
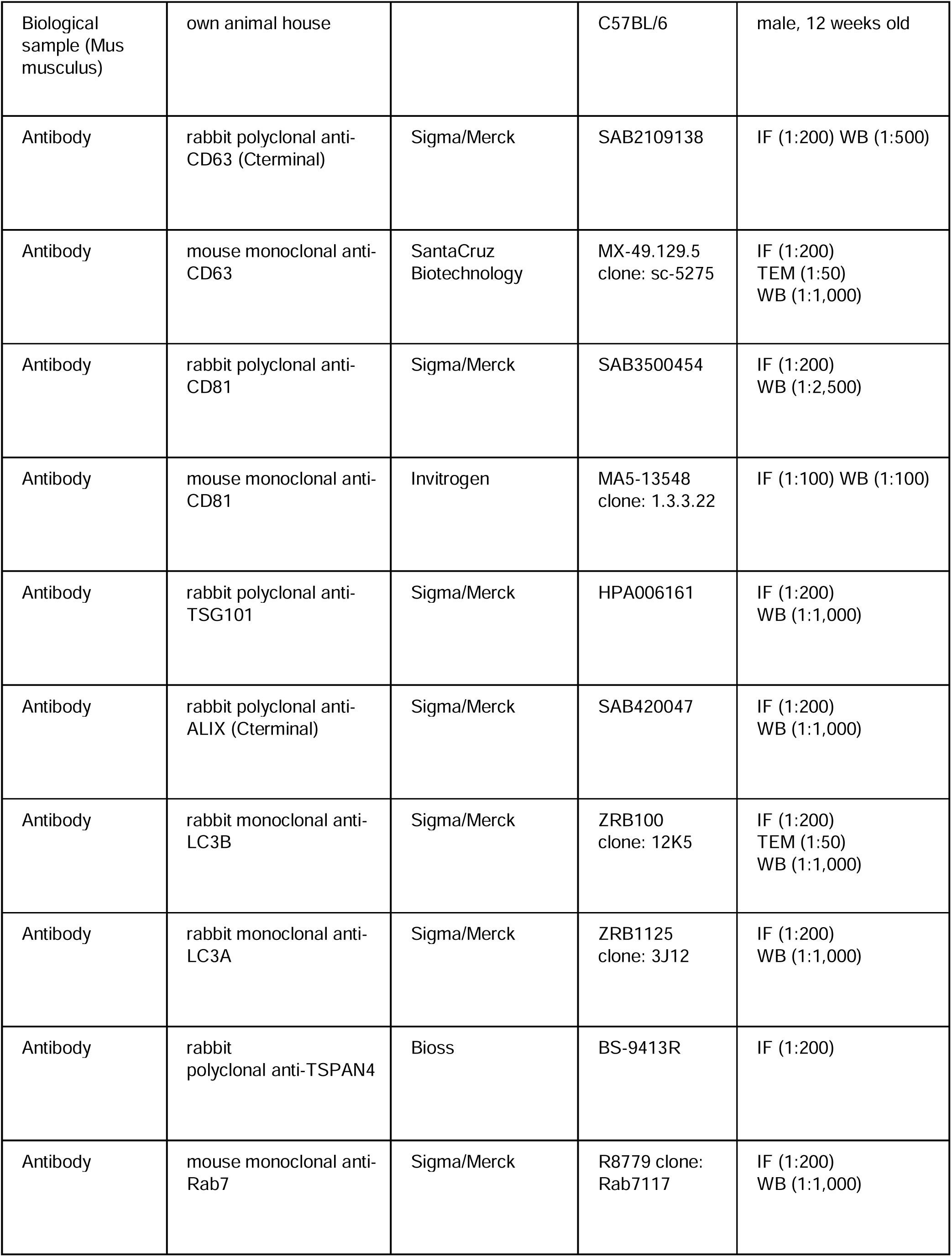
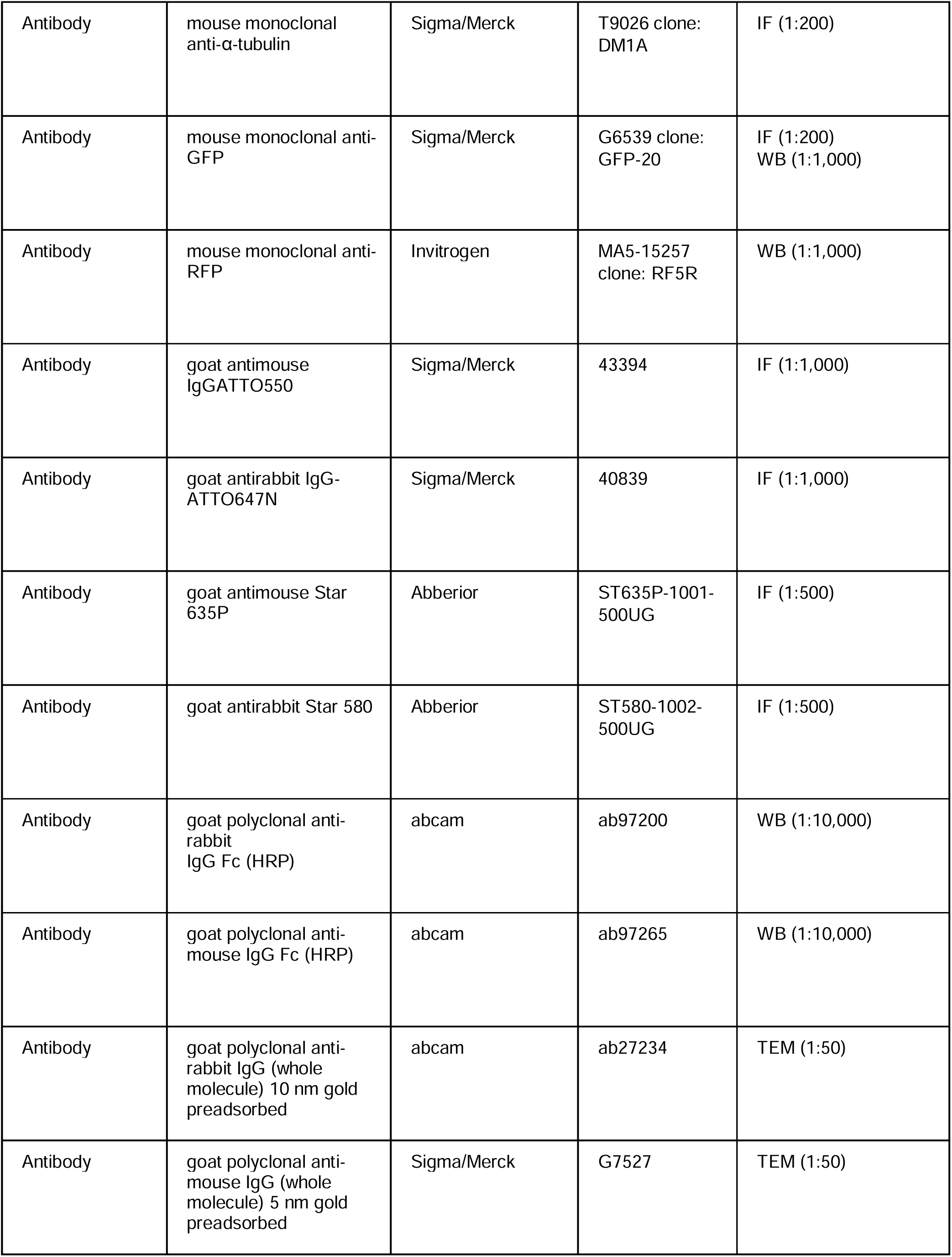
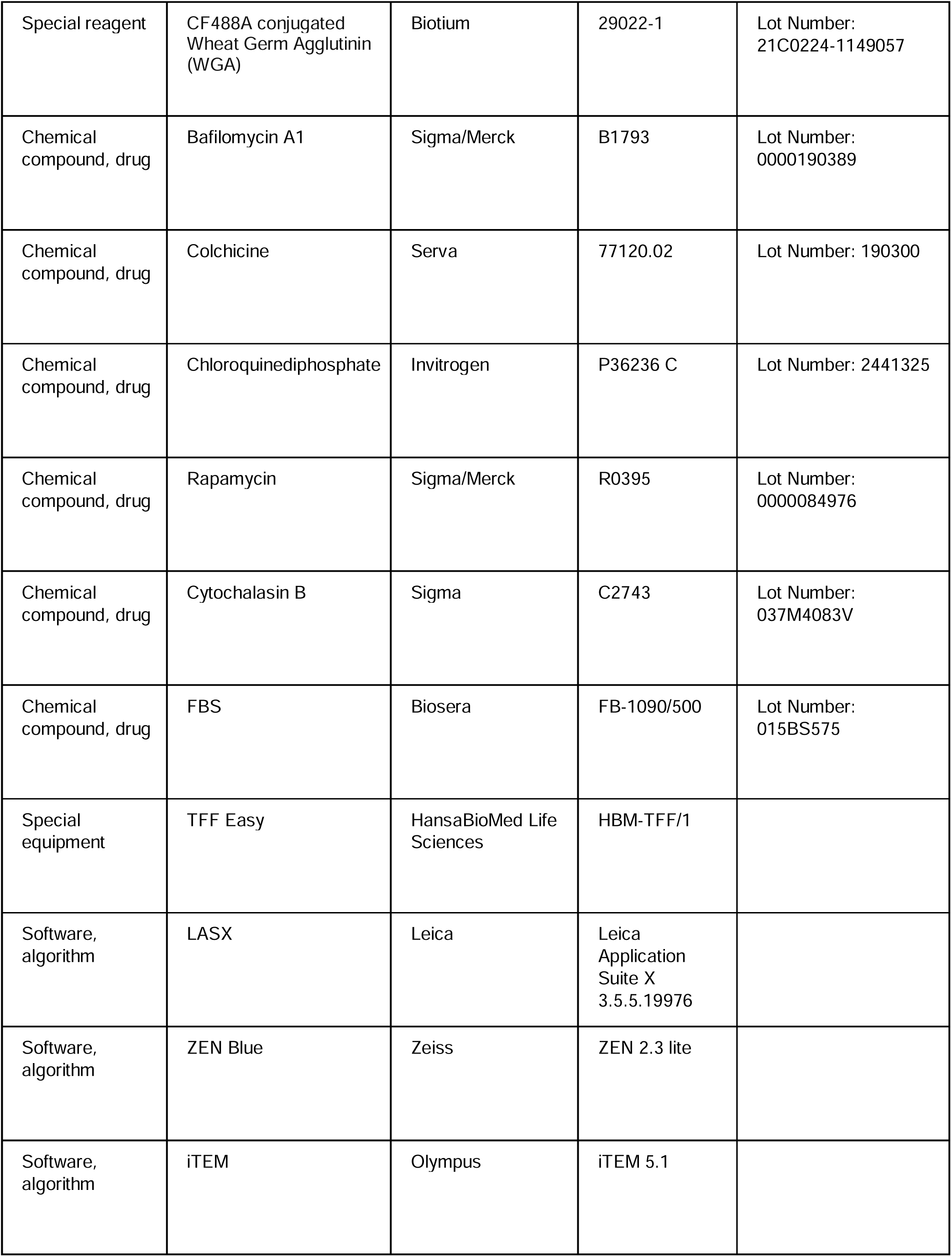
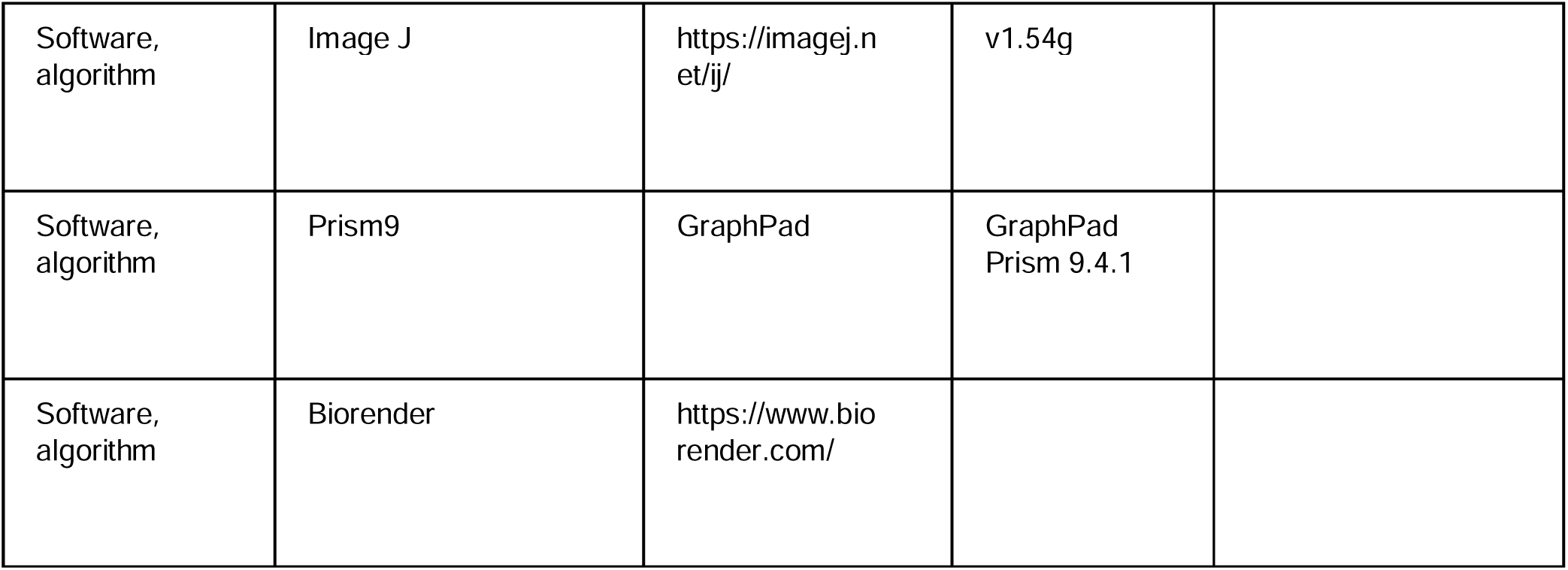

### Cell lines

The HEK293 human embryonic kidney, the HepG2 human hepatocyte carcinoma cell line, the HT29 human colon adenocarcinoma cell lines and the H9c2 rat cardiomyoblast cell line were purchased from the European Collection of Authenticated Cell Cultures (ECACC) through their distributor (Sigma). The HL1 cell line was purchased from Millipore. The HEK293TPalmGFP human embryonic kidney cells was kindly provided by Charles P. Lai^34^. Mouse bone marrow-derived mast cells (BMMCs) were differentiated and expanded as we described previously^35^. The HEK293, HEK293TPalmGFP, HepG2 and H9c2 cell lines were grown in DMEM (Gibco)^36–38^, the HT29 cells were cultured in RPMI 1640 (Gibco)^12^, while the HL1 cells were grown in Claycomb medium^36^. All cells were cultured with 10 % fetal bovine serum (FBS, BioSera) in the presence of 100 U/mL of Penicillin and 100 µg/mL Streptomycin (Sigma). Before analysis by confocal microscopy, the cells were cultured on the surface of gelatin-fibronectin coated glass coverslips (VWR). The coating solution contained 0.02 % gelatin (Sigma) and 5 mg/mL fibronectin (Invitrogen). Coverslips were coated overnight (O/N) at 37 °C.

For transmission electron microscopy (TEM), the adherent cells (HEK293, HEK293T-PalmGFP, HepG2, HT29 and HL1) were grown in gelatin-fibronectin coated 8-well Flux Cell Culture Slides (SPL).

Cell cultures were tested regularly for Mycoplasma infection by PCR, with the following PCR primers: GAAGAWATGCCWTATTTAGAAGATGG and CCRTTTTGACTYTTWCCAC-CMAGTGGTTGTTG ^36^.

### Generation of HEK293T-PalmGFP-LC3RFP cell line

For generation of a stable HEK293T-PalmGFP-LC3RFP cell line, HEK293T-PalmGFP cells were transfected by LentiBrite RFP-LC3 Lentiviral particles (Merck) according to the instructions of the manufacturer. The GFP-RFP double positive cells were sorted by a HS800 Cell Sorter (SONY) and cell banks (master and working cell banks) were prepared. The success of the stable transfection was analyzed by immunocytochemistry and Western blotting. Results are shown in Fig.2_S5.

### Confocal microscopy

Confocal microscopy was carried out as we described earlier^36^ with some modifications. As serum starvation significantly affects autophagy^39^, and EV-depleted FBS in the cell culture medium may influence cellular physiology and morphology^40^, FBS was not removed before fixation. Our study focuses on large EV (lEVs) with diameter > 350-500 nm. EVs in this size range, are neglectibe in FBS because of sterile filtration and heat inactivation of the serum. Unlike the majority of the studies in the field of EVs, here we analyzed untouched, *in situ* fixed and cultured cells together with their microenvironment. Since centrifugation may disrupt the limiting membrane of amphiectosomes, the *in situ* fixation made it possible to observe them in their intact form. The culture medium was gently removed by pipetting from above the cells leaving a thin medium layer only (approximately 150 µL of liquid on the cells). Without any further washing, cells were *in situ* fixed by 4 % paraformaldehyde (PFA) in phosphate buffer saline (PBS) for 20 min at room temperature (RT). The released lEVs were either fixed and captured during the release or were preserved on the gelatin/fibronectin surface coating. After fixation, 3x 5 min washes with 50 mM glycine in PBS were carried out. In the case of the non-fluorescent HepG2 and HT29 cells, a lactadherin-based plasma membrane staining was performed^14,35,37^. Lactadherin (Haematologic Technologies) was conjugated to ATTO488 fluorophore (abcam) according to the instructions of the manufacturer. The lactadherin-ATTO488 conjugate was added to the fixed cells in 1:100 dilution in PBS (for 1 h, RT) before permeabilisation. The unbound lactadherin was removed by washing with PBS (3 times, 5 min, RT) and post-fixation was carried out by 4 % PFA (20 min, RT). PFA was removed by washes with 50 mM glycine in PBS (3 times, 5 min, RT). Blocking and permeabilization of the cells were performed by 10 % FBS with 0.1 % TritonX-100 (Sigma) in PBS (1 h, RT). In general, primary antibodies were applied in 1:200 dilution overnight (O/N) at 4 °C in the above blocking and permeabilization solution. Excess primary antibodies were eliminated by washing with the blocking and permeabilization solution (3 times, 5 min RT). The secondary antibodies were applied in 1:1000 dilution in 1 % FBS in PBS (1 h, RT). Unbound secondary antibodies were eliminated by washing (1 % FBS, in PBS, 2 times, 5 min; PBS 2 times, 5 min; water 2 times, 5 min) and the samples were mounted in ProLong Diamond with DAPI (Invitrogen).

In order to provide evidence for the plasma membrane origin of the outer membrane layer of amphiectosomes, HEK293 were cultured on glass cover slips (VWR). Reaching 60 % confluency, the cells were incubated in expansion medium with 5 µg/mL CF488A conjugated Wheat Germ Agglutinin (WGA) for 30 min at 37°C. After labelling the surface of the plasma membrane by WGA, cells were washed three times by expansion medium and were cultured for an additional 3 hours. Next, they were fixed by 4 % PFA (20 min, RT). LC3B and CD63 labelling were performed as described above.

Microscopic slides were examined by Leica SP8 Lightning confocal microscope with adaptive lightning mode using a HC PL APO CS2 63x/1.40 OIL objective with hybrid detector. Where we showed released MV-lEVs, they were not joined to cells in another detected Z-plane. The applied lookup tables (LUT) were linear during this study. For image analysis and co-localization studies, we applied Leica LASX software using the unprocessed raw images. In case of co-localization studies, 20 % threshold and 10 % background settings were applied.

### Multi-channel STED super-resolution imaging

Immunofluorescent labeling for multi-channel stimulated emission depletion (STED) nanoscopy was performed as in the case of confocal microscopy. The used primary antibodies were: LC3B (rabbit) and CD63 (mouse). Abberior Star 635P Goat anti Mouse Abberior Star 580 Goat anti Rabbit secondary antibodies for STED microscopy have been obtained from Abberior GmbH. Samples were mounted with SlowFade™ Diamond Antifade Mountant (Thermo). Immunofluorescence was analyzed using an Abberior Instruments Facility Line STED Microscope system built on an Olympus IX83 fully motorized inverted microscope base (Olympus), equipped with a ZDC-830 TrueFocus Z-drift compensator system, an IX3-SSU ultrasonic stage, a QUADScan Beam Scanner scanning head, APD detectors and an UPLXAPO60XO 60X oil immersion objective (NA 1.42). We used the 488, 561 and 640 nm solid state lasers for imaging, and a 775 nm solid state laser for STED depletion. Image acquisition was performed using the Imspector data acqusition software (version: 16.3.14278-w2129-win64).

### Purification of sEV fraction

Small EV fractions for TEM analysis were separated from serum-free conditioned medium of HEK293T-PalmGFP cells by gravity-fitration, differencial centrifugation and tangentional flow filtration (TFF Easy, HansaBiomed) as described previosly^38^.

### Transmission electron microscopy

Adherent cells (HEK293, HEK293T-PalmGFP, HepG2, HT29 and HL1), as well as mouse (C57BL/6) kidney and liver tissues pieces (approx. 1.5 mm x 1.5 mm) were immersed in and fixed by 4 % glutaraldehyde (48 h, 4 °C), post-fixed by 1 % osmium tetroxide (2 h, RT) and were embedded into EPON resin (Electron Microscopy Sciences) as described previously^41^. In the case of BMMCs, 920 μL cell suspension was complemented with 80 μL 50 % glutaraldehyde to reach the final 4 % glutaraldehyde concentration. Cells were fixed for 48 h at 4 °C and were post-fixed by 1% osmium tetroxide (2 h, RT). During sample preparation, BMMCs were collected by gravity-based sedimentation. Due to the high viscosity of EPON resin, BMMCs were embedded in LR White low viscosity resin (SPI Supplies) according to the instructions of the manufacturer. Ultrathin sections (60 nm) were contrasted by uranyl acetate (3.75 %, 10 min, RT) and lead citrate (12 min, RT).

For immunogold labelling of ultrathin sections, cells and tissues were fixed by 4 % PFA with 0.1 % glutaraldehyde (48 h, 4 °C) and were post-fixed by 0.5 % osmium tetroxide (30 min, RT). Samples were embedded into LR White hydrophilic resin. The sections were exposed to H_2_O_2_ and NaBH_4_ to render the epitopes accessible and were immunogold labelled as described previously^12^. The contrast was enhanced by uranyl acetate (3.75 %, 1 min, RT) and lead citrate (2 min, RT).

HEK293T-PalmGFP-derived sEVs separated from serum-free conditioned medium, were detected by negative-positive contrasting without embedding and sectioning^15^. Immunogold labelling was performed as described previously^36^. Antibodies were used in 1:50 dilution.

Detailed list of the used antibodies is available in the Key Resources Table.

For all electron microscopic studies, a JEOL 1011 transmission electron microscope was used. Images were captured with the help of Olympus iTEM software and for image analysis, Image J software was used.

### Live cell imaging

The HEK293T-PalmGFP-LC3RFP stable cell line was cultured the same way as HEK293T-PalmGFP cells. Before the experiments, gelatin-fibronectin coated 10-well coverslip bottom chamber slide (Greiner-BioOne) was seeded and treated by 30 μM Chloroquine O/N. Release of migrasomes, amphiectosomes and sEVs were followed by the Leica SP8 Lightning confocal microscope equipped with an Okolab environmental chamber and a Zeiss ELYRA 7 with Lattice SIM² super-resolution fluorescent microscope with the help of 63x/1.4 planapochromat Oil objective. For image analysis, we applied Leica LASX, Zeiss ZEN Blue and Image J softwares.

### Modulation of amphiectosome release

In order to test the release mechanism of amphiectosomes and to distinguish them from migrasomes, different treatments were applied overnight in fresh, serum containing cell culture medium except for Colchicine, where 1 h treatment was selected. Maturation and fusion of endosomes and lysosomes were inhibited by 30 μM Chloroquine (Invitrogen) or 10 nM BafilomycinA1 (Sigma). Actin polymerization was inhibited by 125 ng/mL Cytochalasin B (Sigma). Tubulin polymerization and function were inhibited by 250 pg/mL Colchicine, while an autophagy-related degradation was induced by 50 ng/mL Rapamycin. The selected concentrations were determined based both on literature data and our preliminary experiments (Fig.3_S1). Cellular metabolic activity was determined by a metabolic activity-based Resazurin assay^36^. Fresh cell culture medium was added to control cultures a day before the *in situ* fixation. Reagents were diluted in fresh cell culture medium. Leica TCS SP8 Lightning confocal microscope was used for detection of amphiectosome release. A few hundred µm^2^ sized area with 15-20 µm in height was tile-scanned with a few hundred cells. The MV-lEVs were recognized as CD63 positive EVs surrounded by GFP positive membrane. They were counted and were normalized to the number of nuclei. Raw images were deposited in BioImage Archive (https://www.ebi.ac.uk/bioimage-archive/) with the accession number S-BIAD1456 (doi: 10.6019/S-BIAD1456).

### Western blotting

Presence of proteins and specificity of the used primary antibodies were confirmed by Western blotting as described previously^36^. For accurate quantification (free from variations potentially caused by EV purification), we analyzed cell-, serum- and large EV (diameter > 800 nm) free conditioned medium. The cells were cultured O/N in a serum-free culture medium. After harvesting cells were eliminated by centrifugation (300 g, 10 min at 4 °C) followed by a 2.000 g centrifugation (30 min at 4 °C) to eliminate lEVs. Total protein content of the conditioned, serum-, cell- and lEV-free medium was precipitated by trichloroacetic acid as described previously^36,42^. The protein pellets were suspended in cOmplete Protease Inhibitor Cocktail (Roche) containing radio-immunoprecipitation assay (RIPA) buffer.

When whole cell lysate was tested for validation of antibodies and the HEK293T-PalmGFP-LC3RFP cell line, cells were lysed in cOmplete Protease Inhibitor Cocktail (Roche) containing RIPA buffer.

Polyacrylamide gel electrophoresis was carried out using 10 % gels (acrylamide/bis-acrylamide ratio 37.5:1) or any kDa precast gels (Biorad) and a MiniProtean (BioRad) gel running system. For better solubilization of membrane proteins, equal volumes of 0.1 % TritonX-100, Laemmli buffer and samples were mixed as described previously^43^. Approximately 10-30 µg protein were loaded into each well. Following electrophoretic separation, proteins were transferred to PVDF membranes (Serva). Membranes were blocked with 5 % skimmed milk powder or 5 % BSA in washing buffer for 1 h. Primary antibodies were applied in 1:1000 dilution with the exception of the anti-CD63 (rabbit), anti-CD81 (rabbit) and anti-CD81 (mouse) antibodies where 1:500, 1:2500 and 1:100 dilutions were used, respectively. Peroxidase-labelled secondary antibodies were applied in 1:10 000 dilution. The signals were detected by ECL Western Blotting Substrate (Thermo Scientific) with an Imager CHEMI Premium (VWR) image analyzer system. In case of quantification, equal protein amounts were loaded to the gels. Within a biological replicate, the control and Chloroquine-treated samples were run on the same gels. To enable comparison, the relative expression of control and Chloroquine-treated samples were determined and compared.

### Software and statistical analysis

For image capturing, analysis and co-localization studies, Leica LAS X, Zeiss ZEN Blue, Olympus iTEM and Image J software were used. Figures and graphs were generated using GraphPad Prism 9.4.1 and Biorender (BioRender.com). For statistical analysis, standard deviation was calculated. Unpaired two tailed Student’s t-tests and one-way ANOVA were used (* p<0.05, ** p<0.01, *** p<0.001, **** p<0.0001).

## Acknowledgements

This research was funded by the NVKP_16-1-2016-0004 grant of the Hungarian National Research, Development and Innovation Office (NKFIH), ÚNKP-23-3-I-SE-2, the Semmelweis Innovation Fund (STIA 2020 KFI), Hungarian Scientific Research Fund (OTKA K120237 and FK 138851), Eötvös Loránd University Excellence Fund (EKA 2022/045-P101), Hungary Academy of Sciences, LP2022-13/2022, VEKOP-2.3.2-162016-00002, VEKOP-2.3.3-15-2017-00016, the Higher Education Excellence Program (FIKP) and the Therapeutic Thematic Programme TKP2021-EGA-23. This study was also supported by the grants RRF-2.3.121-2022-00003 (National Cardiovascular Laboratory Program) and 2019-2.1.7-ERA-NET-2021-00015. The project has received funding from the EU’s Horizon 2020 Research and Innovation Programme under grant agreement No. 739593, „Momentum” research grant from the Hungarian Academy of Sciences (LP2022-5/2022) and the Hungarian Brain Research Program NAP2022-I-1/2022. TV and CC was supported by the János Bolyai Research Scholarship of the Hungarian Academy of Sciences.

The authors would like to thank the ZEISS Microscopy Customer Center Team for the collaboration, and Dr. Abel Pereira da Graça for the possibility to use the Elyra7 SIM^2^ super-resolution microscope. The authors are grateful to Györgyné Vidra, Györgyi Balogh and Andrea Orbán for their technical help and advises.

