## Supplementary Material for "A “torn bag mechanism” of small extracellular vesicle release via limiting membrane rupture of *en bloc* released amphisomes (amphiectosomes)"

### Fig.1_S1. Additional Transmission electron micrographs of mouse kidney and liver sections

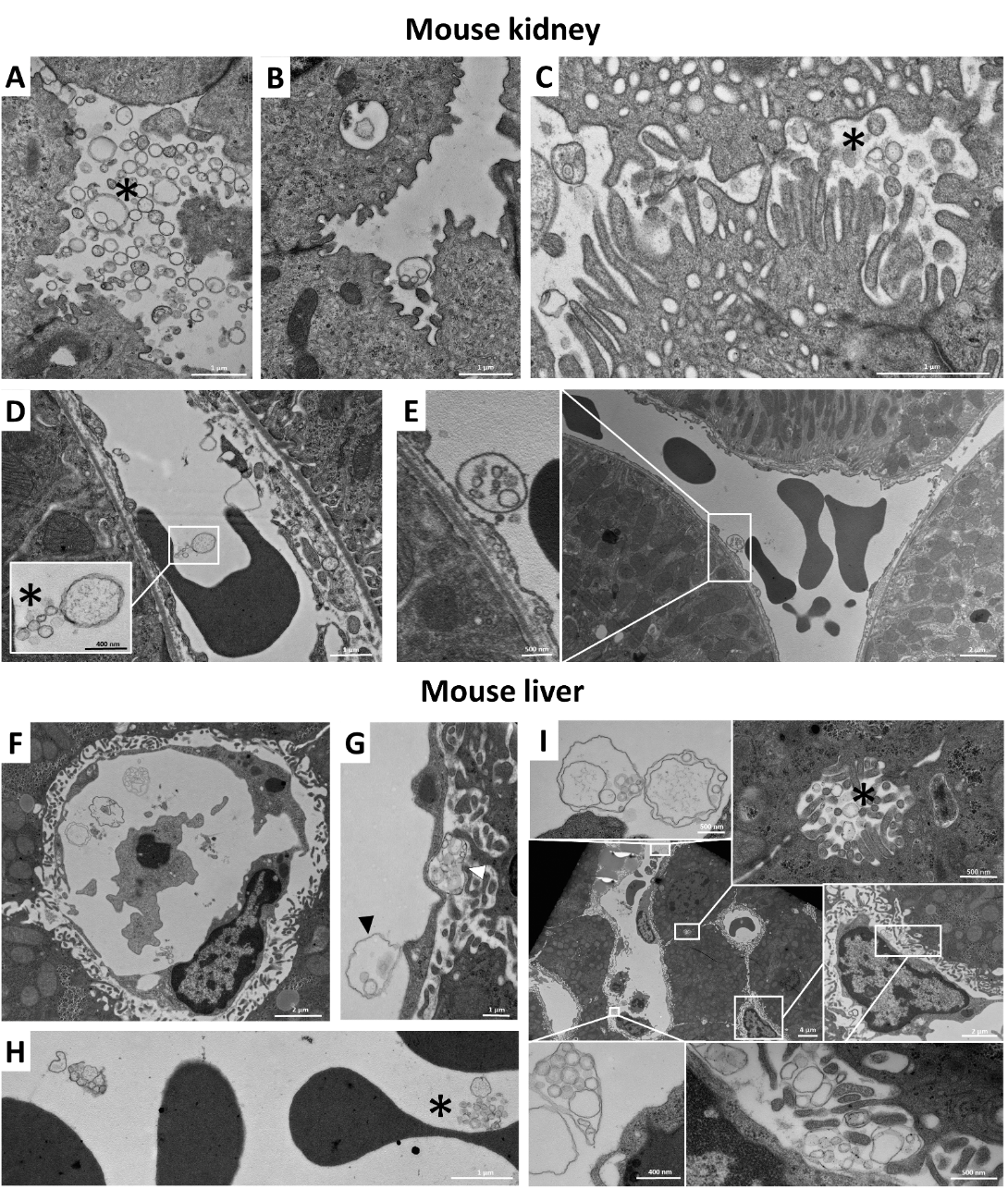

For further evaluation of the presence of different EVs in mouse kidney (A-E) and liver (F-I), low and high magnification images are shown. Presence of invidual small and large EVs is indicated by asterix in the case of mouse kidney (A, C, D) and mouse liver (F, H, I). The MV-lEVs and individual EVs were present simultaneously (C, D, F, H, I). In the mouse liver ultrathin section (G), MV-lEV secretion by endothelial and subendothelial cells (black and white arrow heads, respectively) were detectable in the same image.

### Fig.2_S1. Localisation of GFP signal in HEK293T-PalmGFP cells

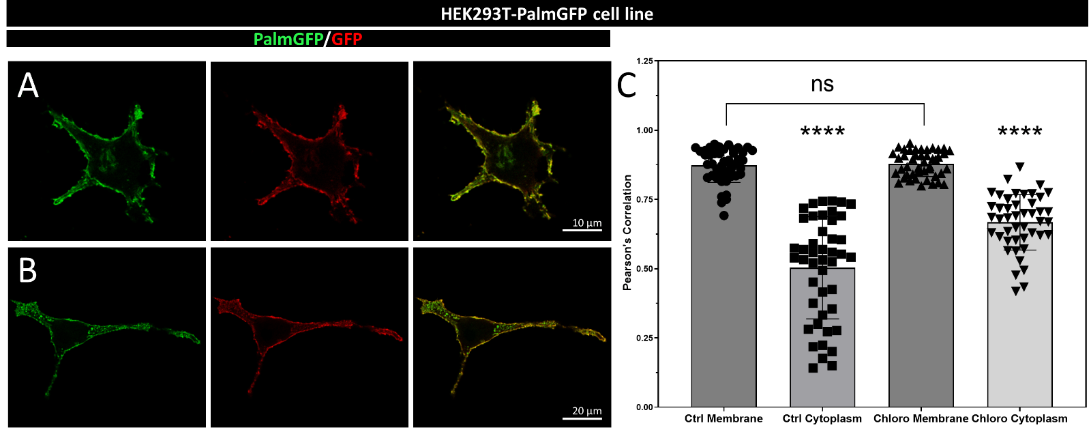

The GFP positivity of the HEK293TPalmGFP cells was controlled by immunofluorescence microscopy on control (A) and 30 µM Chloroquine-treated (B) cells. Co-localization of Palm-GFP and the anti-GFP signal was calculated in the plasma membrane and in the cytoplasm (C) and Pearson’s correlations were visualized. The green fluorescence in the plasma membrane was clearly GFP-dependent in the case of control (Ctrl Membrane) and Chloroquine-treated (Chloro Membrane) samples, while in the cytoplasm, the correlation was significantly weaker (Ctrl Cytoplasm and Chloro Cytoplasm). Within the cytoplasm, the correlation was stronger in Chloroquine-treated cells, suggesting that endosomal membrane may contain Palm-GFP possibly as a result of the membrane endocytotic or recycling processes (p < 0.0001, t-test; n = 45).

### Fig.2_S2. Confocal microscopic images of amphiectosome release by HT29 and HepG2 cells

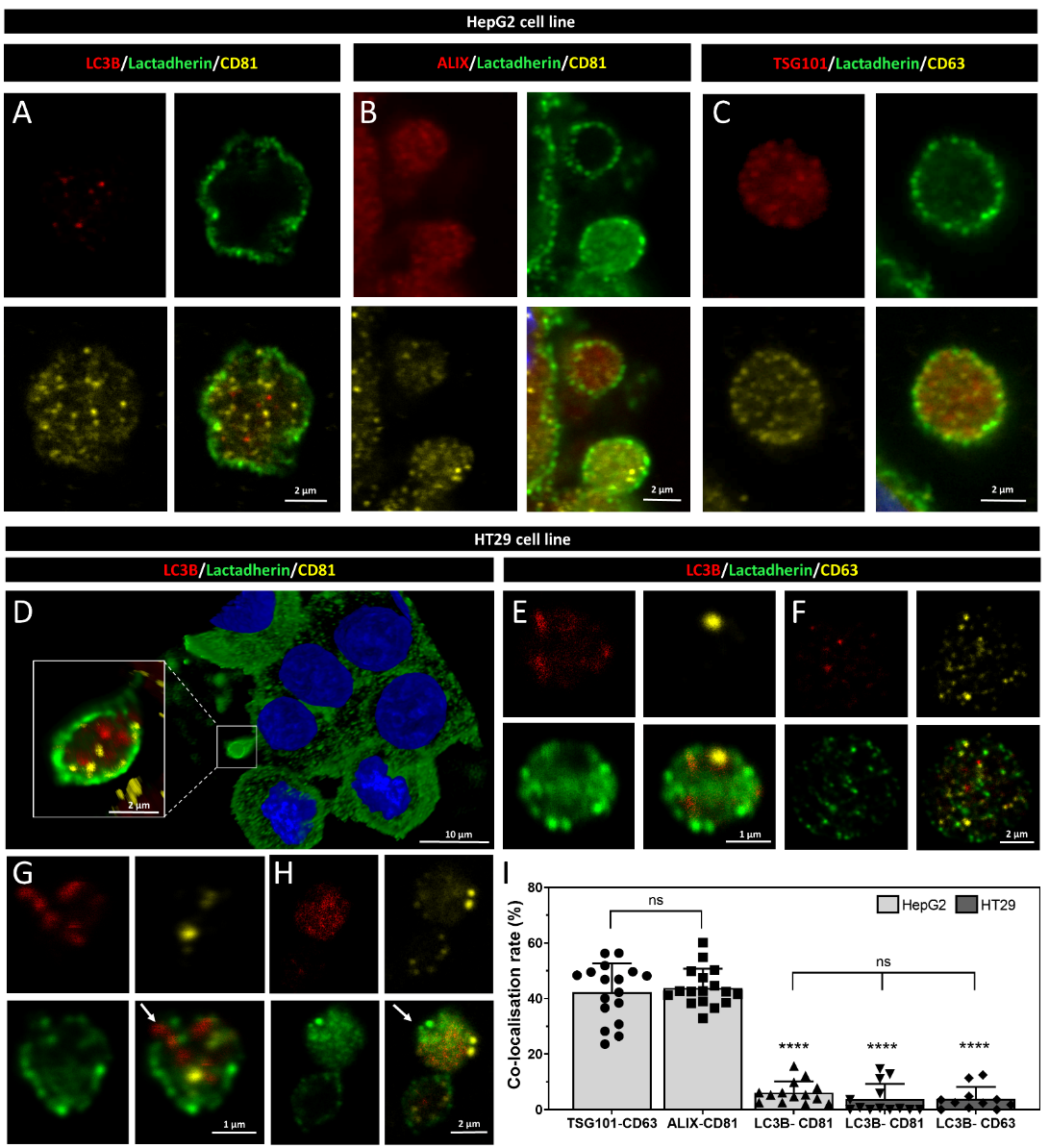

The intraluminal EVs of MV-lEVs were found to be positive for LC3B and CD81 in HepG2 (A) and HT29 cells (D). The presence of ALIX/CD81 (B) and TSG101/CD63 (C) was also examined in the released amphiectosomes of the HepG2 cell line. In the case of HT29 cells, phases of the "torn bag mechanism” were captured, including a secreted intact amphiectosome (E), amphiectosomes with ruptured limiting membrane releasing internal vesicles (G, white arrow), an inside-out secreted amphiectosome (H, white arrow), and an amphiectosome with a fully disintegrated limiting membrane and released sEVs (F). Co-localization rates of marker proteins were calculated (I). The typical sEV markers of HepG2 co-localized with eachother. Significant difference was not found among them. In contrast, low co-localization rates were detected between the “classical” sEV markers (CD63 and CD81) and LC3B. The co-locolasation rates between “classical sEV markers - LC3B and within “classical” EV markers (TSG101 - CD63 and ALIX - CD81) were found significaltly different (one-way ANOVA, p<0.0001, n=11-17 confocal images).

### Fig.2_S3. Additional confocal microscopic images of H9c2, HEK293T-PalmGFP and HepG2 cells derived MV-lEVs

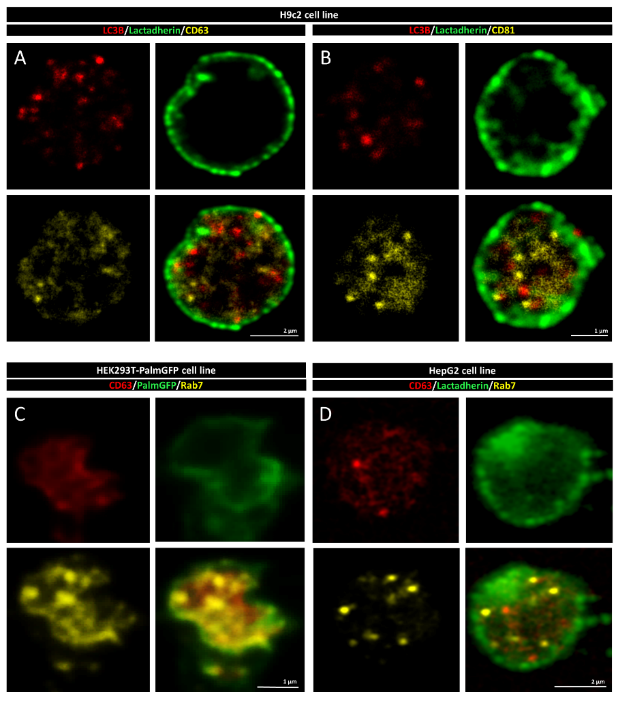

Released amphiectosomes of H9c2 rat cardiomyoblast cell line were captured. They contain CD63, CD81 or LC3B positive ILVs (A,B). The MV-lEVs released by HEK293T-PalmGFP (C) and HepG2 cells (D) were tested for CD63 and the Rab7 late endosomal marker. Both CD63 and Rab7 were present in association with the ILVs.

**Fig.2_S4. Qualitative Western blot validation of antibodies used in immunofluorescence detection**

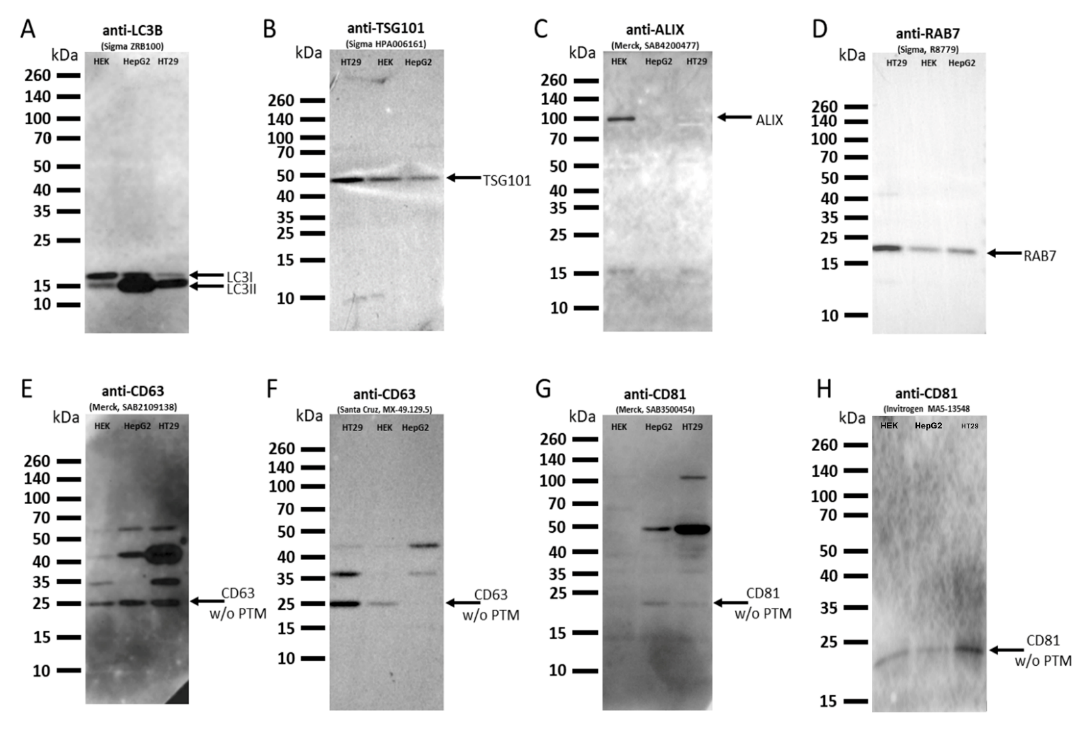

Whole cell lysates from three distinct human cell lines (HEK293, HepG2 and HT29) were employed in the validation process to reduce the cell line-specific variations in this qualitative study. Protein bands lacking posttranslational modifications are noted as "w/o PTM." The found posttranslational modifications of CD63 and CD81 are widely and well recognized, the results are in agreement with the Western blot data provided by the antibody suppliers.

**Fig.2_S5. Characterization of the in house generated HEK293T-PalmGFP-LC3RFP cell line.**

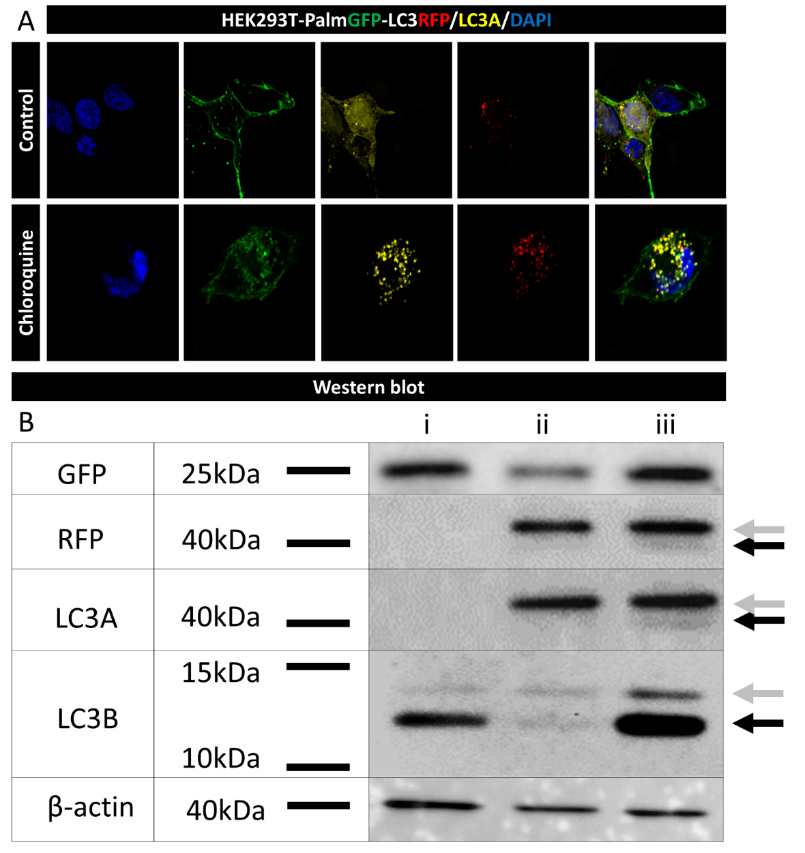

Confocal microscopy (A) and Western blot analysis (B) were employed to characterize the HEK293T-PalmGFP-LC3RFP cell line. For overnight Chloroquine treatment, 30 μM Chloroquine was applied. Punctate LC3 fluorescence observed in (A) corresponds to autophagosomes or amphisomes. In panel (B), whole-cell lysates from HEK293T-PalmGFP (lane i), HEK293T-PalmGFP-LC3RFP (lane ii) and Chloroquine-treated HEK293T-PalmGFP-LC3RFP (lane iii) samples were examined. Gray arrows indicate LC3I, while black arrows highlight the lipidated LC3II. Full Western Blot images can be find in Fig.3_S2B supplementary figure.

### Fig.2_S6. Structures involved in amphiectosome release

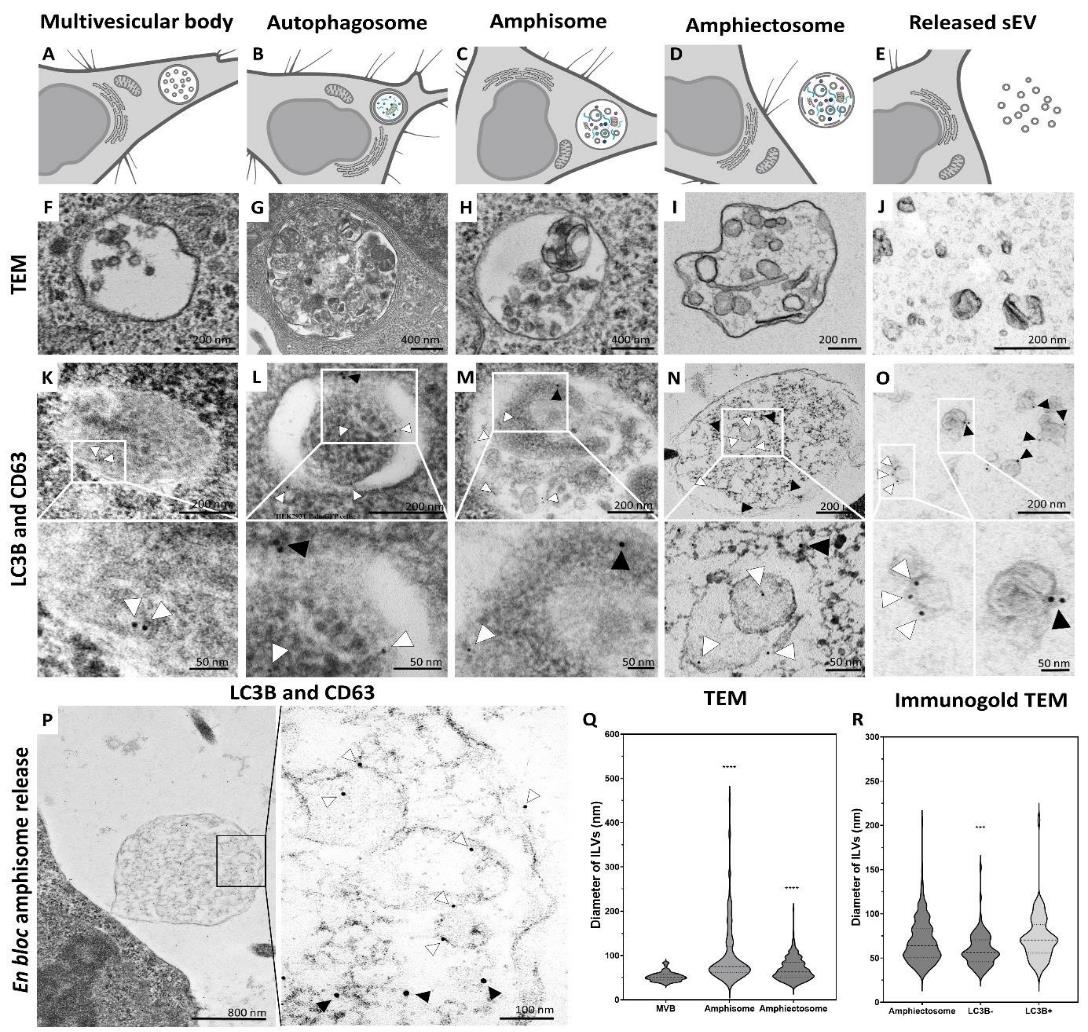

Multivesicular body (MVB, A), autophagosome (B), amphisome (C), amphiectosome (D) and secreted sEVs (E) were identified by TEM with and without immunogold labelling of HEK293T-PalmGFP cell cultures. White arrowheads (5 nm gold particles) indicate CD63 and black arrowheads (10 nm gold particles) show LC3B. While MVBs (F) were LC3B negative (K), we detected CD63 positivity on the surface of the ILVs (K). In an autophagosome (G), the limiting membrane layers were positive for CD63 and LC3B (L). In contrast, the internal membranes of autophagosome were CD63 single positive (L). In the case of an amphisome (H), heterogeneous membrane structures were visible with variable size and morphology. The ILVs were either CD63 or LC3B positive (M). The amphiectosomes were located in the extracellular space and contained ILVs of different size and shape (I). The ILVs of amphiectosome (as in case of amphisome), were either CD63 or LC3B positive (N). Secreted sEVs purified from serum-free conditioned medium (J) with immunogold labelling (O), were also found to be either CD63 or LC3B positive. Release of an amphiectosome is shown (P) with CD63 and LC3B immunogold signals. Higher magnification of the insert is indicated by the black rectangle. It shows either CD63 or LC3B positive ILVs. Size distributions of ILVs of MVBs, amphisomes and amphiectosomes were determined on Epon-embedded ultrathin sections (Q). Although the ILV sizes differed significantly (one-way ANOVA, ****: p<0.0001, n=73, 138 and 595, respectively), the majority of ILVs had a diameter between 40-100 nm. The diameter of LC3B positive and negative ILVs of amphiectosomes was assessed on TEM images of immunogold labelled ultrathin sections (R). LC3B negative ILVs were significantly smaller than the LC3B positive ones, while the ILVs in the Epon embedded sections did not differ from the LC3B positive ones (one-way ANOVA, p <0.001, n= 595, 101 and 70, respectively).

### Fig.3_S1. Supporting information for treatments and size distribution of MV-lEVs

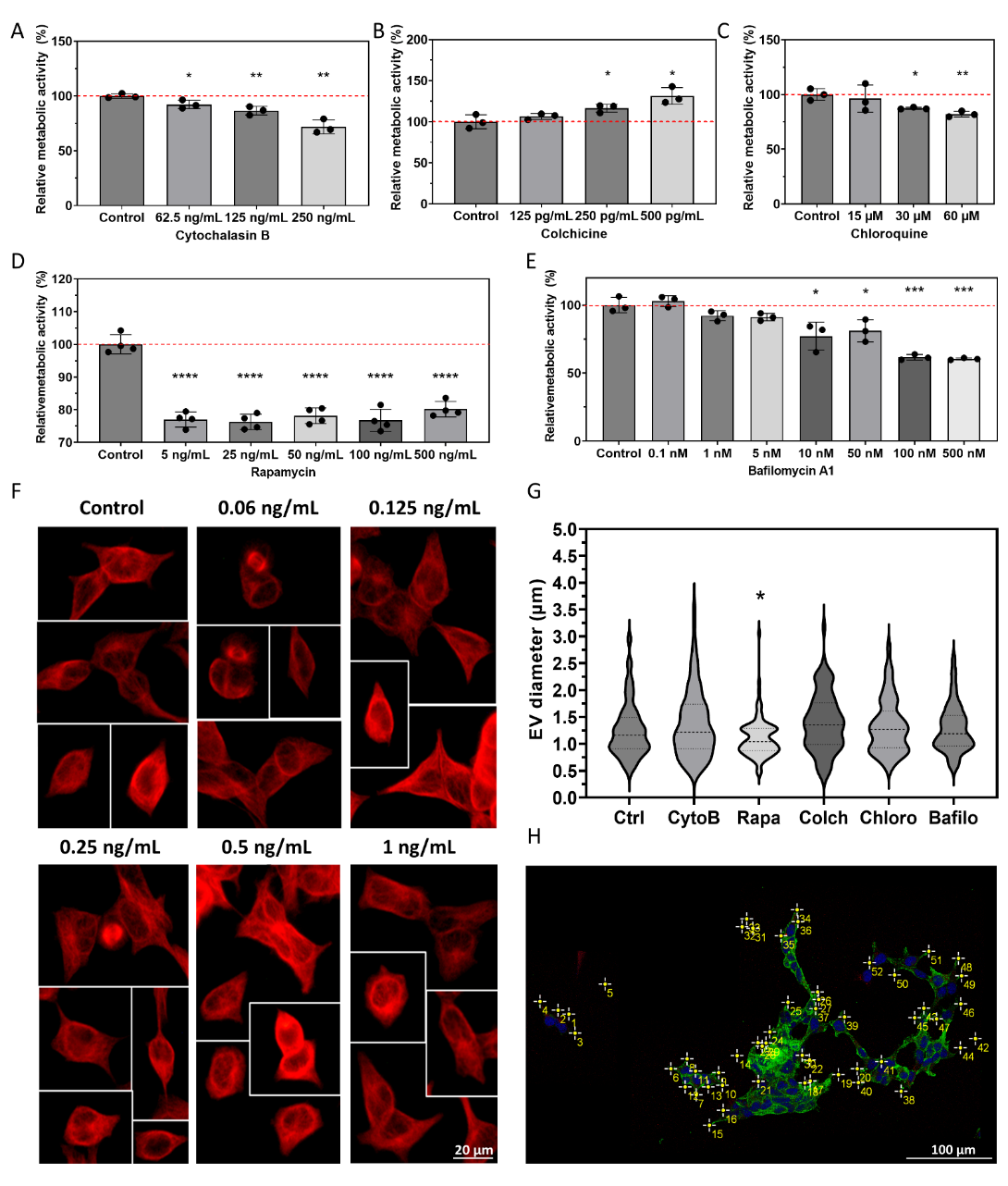

Relative metabolic activity was assessed through a Resazurin assay (A-E) during treatment optimization. The red dashed line indicates 100 % metabolic activity, representing control cells. Results are presented as mean ± SD values of n=3-4 independent biological replicates. Student’s unpaired t-test was performed to compare control and treated cells (*: p<0.05, **: p<0.01, ****: p<0.0001). For Colchicine treatment, alterations in the microtubular network were observed through immunocytochemistry (F) and documented using an epifluorescent microscope. Changes in the diameter of released large multivesicular EVs (MV-lEVs) under various treatments were determined on confocal microscopy images. A significant reduction in size was identified only in the case of Rapamycin treatment (G, one-way ANOVA test *: p<0.05, n= 95-101). An example for MV-lEV number calculation based on confocal images is shown (H).

### Fig.3_S2. Unedited Western blots of Fig.2_S4 and Fig.3

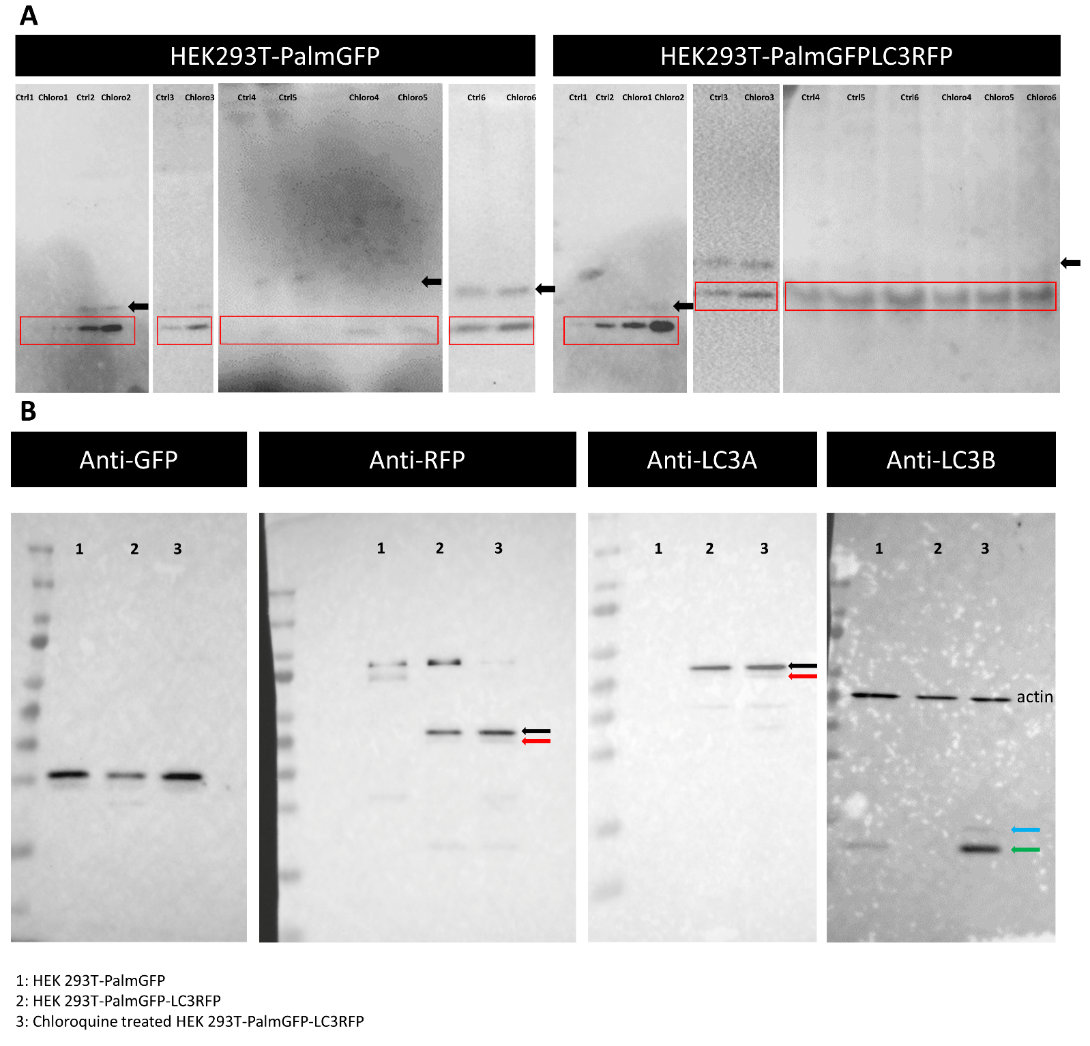

Subfigure A displays the unedited Western blots utilized for the quantification in Fig.3J. Black arrows indicate the location of LC3I, while red squares represent the quantified LC3II. In Subfigure B, the unedited blots from Fig.2_S5B are presented. The PVDF membrane was cut into four pieces after blotting before immunological testing. Black arrows denote the LC3I-RFP fusion protein, while red arrows indicate the LC3II-RFP fusion protein. The blue arrow points to the native LC3I, while the green arrow represents the native LC3II protein.
